# Sleep complaints in alexithymia reflect non-specific negative affectivity

**DOI:** 10.64898/2026.07.29.741441

**Authors:** Pardis Adibi, Péter Przemyslaw Ujma

## Abstract

Alexithymia is the inability to identify and describe emotions, associated with an increased risk of somatic and psychiatric disease. Alexithymia is also characterized by reduced subjective sleep quality. However, it is not well-known if this reduction in sleep quality reflects physiological changes in sleep, a specific alteration in perceived sleep quality not captured by objective metrics, or a non-specific spillover of the negative affective style which characterizes alexithymia. In BSETS, a large (N=228) EEG study of naturalistic sleep, we found that alexithymia symptoms are associated with worse habitual (AIS) and current (GSQS) subjective sleep quality (β_AIS_=0.207, p_AIS_=0.002, β_GSQS_=0.203, p_GSQS_=0.002). However, we observed no significant association between alexithymia and EEG-based sleep efficiency (β=- 0.017, p=0.791). The association with subjective sleep quality was almost completely attenuated when controlling for either depressive symptoms (PHQ-9, β_AIS_=0.001, p_AIS_=0.982, β_GSQS_=0.044, p_GSQS_=0.486) or trait neuroticism (ZKPQ, BFI-44, β_AIS_=0.057, p_AIS_=0.442, β_GSQS_=0.048, p_GSQS_=0.514). A possible exception is the Difficulty Identifying Feelings alexithymia dimension, reflecting complaints about the labeling of interoceptive experiences, which was associated with lower subjective sleep quality even after covariate control. This pattern of findings suggests that alexithymia is characterized only by alterations of subjective sleep quality, and even these reflect a non-specific increase in negative affect which influences self-reports of sleep quality, rather than a specific reduction in perceived sleep quality itself.

## Introduction

Alexithymia is a difficulty in identifying and describing emotions (Hogeveen and Grafman, 2021; Sifneos, 1973), including a difficulty identifying the physical arousal associated with feelings, limited imagination, and an externally oriented thinking style. Alexithymia is generally measured as a continuous trait, ranging from no symptoms through the subclinical range to clinically significant symptoms using the Toronto Alexithymia Scale (Bagby et al., 1994). Alexithymia is a known risk factor of both psychiatric (Ricciardi et al., 2015) and somatic (Taylor et al., 1999) illness, including sleep disorders (Ma et al., 2020).

Sleep problems in alexithymia can be explained by at least three distinct hypotheses. First, it is possible that the functional and structural changes in the brain which characterize alexithymia are also implicated in sleep-related processes, for example, via disruptions of REM regulation (Godin et al., 2013). This hypothesis would predict that alexithymia is intimately linked with sleep physiology and associated with alterations in objectively measured sleep, especially sleep elements implicated in emotional processing. Support for this hypothesis is inconclusive, because studies using objectively measured sleep in alexithymia are rare and typically based on small samples. Bazydlo et al (Bazydlo et al., 2001) in a sample of 50 participants found shallower non-rapid eye movement (NREM) sleep and more fragmented rapid eye movement (REM) sleep at higher levels of alexithymia, an association that survived correction for depressed affect. These findings, however, were not replicated in a smaller study by De Gennaro et al (De Gennaro et al., 2002), who found no significant associations. Bileviciute-Ljungar and Friberg (Bileviciute-Ljungar and Friberg, 2020) in a small sample of myalgic encephalomyelitis patients found no significant associations between TAS scores and sleep. Godin et al (Godin et al., 2013) found higher alexithymia scores in patients with REM sleep behavior disorder, a disorder of the sleep stage with the greatest putative role in emotional processing (Tempesta et al. 2018), but did not investigate whether the degree of alterations of objective sleep correlates with the severity of alexithymia.

Second, it is possible that while objective alterations of sleep are absent, there is a specific deficit in subjectively perceived sleep quality. Although poor subjective sleep quality is a clinically important indicator with independent epidemiological relevance (Utsumi et al., 2022), it is only moderately correlated with objectively measured sleep quality, indicated by sleep efficiency, duration, or fragmentation (Pierson-Bartel and Ujma, 2024; Svetnik et al., 2020). Thus, it is possible that alexithymia either impacts sleep in more subtle ways than what standard polysomnography measures capture, or that it doesn’t impact the physiological character of sleep, only the perception of its quality. This hypothesis predicts no alterations in objective sleep, but lower self-reported sleep quality. Importantly, under this hypothesis lower self-reported sleep quality would be statistically independent from non-specific factors such as neuroticism, anxiety, or depression, which are also elevated in alexithymia (Hendryx et al., 1991; Marchesi et al., 2000) and predict lower self-reported sleep quality (Catherman et al., 2023).

Third, it is possible that alexithymia is not associated with specific alterations in sleep. Lower self-reported sleep quality may only appear because alexithymia is associated with a general pattern of negative emotionality, which impacts subjective reports of sleep as well. In other words, under this hypothesis, sleep quality would be perceived as no more negative than other self-reported characteristics of the patients’life, implying that low subjective sleep quality reflects non-specific affective symptoms or a generally negative response style, and not a specific change in perceived sleep quality itself. This hypothesis predicts that even if alexithymia is associated with low subjective sleep quality, non-specific measures of negative affectivity will fully account for this association.

The literature strongly supports that alexithymia is associated with a reduced quality of self-reported sleep. A recent meta-analysis (Alimoradi et al., 2022) based on 24 studies with 7546 participants found an overall correlation of r=0.44 between sleep quality and alexithymia symptoms, with substantial between-study heterogeneity, but no evidence for publication bias. It is, however, unclear whether this finding supports the second hypothesis of a specific alteration of sleep perception, or the third hypothesis about confounding by non-specific symptoms. At least one large study (De Gennaro et al., 2004) found that the correlation between alexithymia symptoms and subjective sleep quality is fully attenuated after statistically controlling for symptoms of depression and/or anxiety, however, smaller studies (Bazydlo et al., 2001; Han et al., 2023) reported incomplete attenuation.

In sum, it remains unanswered whether alexithymia is associated with objective or only subjective changes in sleep quality, and whether the well-documented changes in subjective sleep quality reflect a specific alteration of sleep, or non-specific affective symptoms. In the present study, we sought to explore the nature of sleep alterations in alexithymia using a large sample of participants with TAS-20 scores, EEG-based objective sleep measures and two different indicators of subjective sleep quality and three indicators of negative affectivity.

## Methods

### Participants

We used data from the Budapest Sleep, Experiences and Traits Study (BSETS). In the current paper we focus on aspects of BSETS relevant to the study at hand; the full protocol has been published separately (Taji et al., 2023).

BSETS is a multiday observational study of the interplay between sleep, trait-level characteristics, and daily experiences of healthy volunteers. BSETS participants were recruited via branched diffusive convenience sampling. Inclusion criteria were minimal to ensure ecological validity and consisted only of an age of at least 18 years, being fluent in Hungarian, and non-recruitment from clinical settings. 267 participants provided at least some data for BSETS: of these, the current study uses 228 with full data on all variables used in primary analyses.

BSETS included a psychometric battery containing validated self-report measures of alexithymia, personality, the quality of habitual sleep, and depressive affect. It also included a tracking period of at least one week, during which participants recorded nightly objective sleep using a Dreem2 mobile EEG device, and self-reported the quality of recent sleep each morning.

The Institutional Review Board (IRB) of Semmelweis University, as well as the Hungarian Medical Council (under 7040-7/2021/ EÜIG _“_Vonások és napi események hatása az alvási EEG-re_”_ [The effect of traits and daily activities and experiences on the sleep EEG]), approved BSETS as compliant with the latest revision of the Declaration of Helsinki. All participants gave written informed consent on a form reviewed and approved by the IRB.

### Psychometric battery

BSETS included a psychometric battery which was used to calculate some variables of interest.

Participants self-reported the quality of their habitual sleep using the Hungarian version (Ronai et al., 2017) of the Athens Insomnia Scale (AIS). In the AIS, participants self-report the degree and frequency of recently occurring sleep problems using a Likert-scale.

Alexithymia was assessed with the Hungarian version of the Toronto Alexithymia Scale (TAS-20) (Cserjési et al., 2007). This questionnaire uses Likert-type questions to assess three dimensions of alexithymia symptoms (difficulty identifying feelings, difficulty describing feelings, and externally oriented thinking), as well as a total alexithymia score. In the primary analysis we used the total TAS-20 score. In secondary analyses, we used TAS-20 subscales Difficulty Identifying Feelings (DIF), Difficulty Describing Feelings (DDF), and Externally Oriented Thinking (EOT).

Depressive symptoms were assessed with the Hungarian version of the Patient Health Questionnaire (PHQ-9) (Kroenke et al., 2001). In this brief screening questionnaire, participants self-report the frequency of experiencing various DSM-IV-based symptoms of depression.

Neuroticism/negative emotionality were assessed using the Emotional Instability dimension of the Hungarian version (Rózsa, 2010) of the Big Five Inventory (BFI-44), and the Neuroticism-Anxiety dimension of the Zuckermann-Kuhlmann Personality Questionnaire (ZKPQ) (Kövi et al., 2018). In both questionnaires, participants self-report characteristics and behaviors associated with these personality traits on either a Likert scale (BFI-44), or as a binary response (ZKPQ).

### Daily sleep measures

BSETS also included a tracking period of at least one week during which participants recorded the quality of their sleep during the previous night and wore an EEG headband each night. Individual averages were calculated from all objective and subjective sleep reports of the tracking period to ensure a single data point for each participant.

Subjective sleep quality of the previous night was self-reported using the Hungarian version of the Groningen Sleep Quality Scale (GSQS) (Simor et al., 2009). The GSQS uses 14 Likert-type items to assess the degree of sleep problems during the previous night.

To assess objective sleep, participants wore a Dreem2 headband each night. Dreem2 uses dry silicone electrodes at positions Fp1, F7, F8, O1 and O2 and records EEG with a sampling frequency of 250 Hz (Dreem Inc, 2017; Taji et al., 2023). Sleep is scored in 30-second segments using a validated algorithm (Arnal et al., 2020), and estimates of sleep onset latency, wake after sleep onset, total sleep time, sleep efficiency, and the duration of percentage of sleep stages N1, N2, SWS, and REM are provided each night. Of these, we operationalized objective sleep quality as sleep efficiency, the percentage of bedtime spent asleep, and used it as the key variable in primary analyses. This was motivated by previous work showing that this metric shows the highest correspondence with subjective sleep quality (Pierson-Bartel and Ujma, 2024; Svetnik et al., 2020). However, in secondary analyses, we also explored the association of other objective sleep measures (total sleep time, wake after sleep onset, sleep onset latency, REM latency, REM, N1, N2, N3 percentage, and the number of awakenings) with alexithymia.

### Statistical analysis

In our study, we calculated the association of both objective and subjective sleep quality with alexithymia symptoms and explored the effect of including depressive symptoms and two measures of neuroticism/emotional instability as confounding variables.

Because sleep efficiency, PHQ and AIS scores had considerable skewness and kurtosis, they were transformed. Sleep efficiency was reflected and log-transformed (SE_new_=log(1-SE)), while PHQ and AIS scores were log-transformed. **Supplementary figure S1** shows the histograms of all transformed and untransformed variables, as well as their correlations and scatterplots.

Internal reliability of questionnaires was calculated using Cronbach’s alpha and McDonald’s omega. For neuroticism-anxiety, based on binary ZKPQ items, McDonald’ s omega was calculated using tetrachoric correlations. For the multidimensional TAS, the same metric is reported as total omega from a hierarchical model with three second-order factors corresponding to the three alexithymia dimensions. All other cases (including GSQS, which, while originally binary, yielded a continuous measure due to averaging results from multiple nights) used a single-factor model based on Pearson correlations.

In primary analyses, we used sleep quality (GSQS scores, AIS scores, and sleep efficiency) as the outcome and TAS-20 total score, age, and sex as predictors. In two additional models, we included controls for either PHQ-9 total scores (depression-controlled models), or ZKPQ Neuroticism-anxiety and BFI Emotional instability scores (personality-controlled models). In secondary analyses, we replaced TAS-20 total scores with dimension scores (DDF, DIF, and EOT), or sleep efficiency with other objective sleep parameters, with a similar inclusion of controls. We adjusted for multiple comparisons using the Benjamini-Hochberg procedure, using separate pools of p-values for the primary and the two secondary analyses.

All data and code needed to replicate our analyses is available at osf.io/j3ntp.

## Results

### Descriptive statistics

Table 1 reports descriptive statistics from the main study variables. All statistics were calculated from a subset of the original BSETS sample with complete cases for all variables (N=228).

Due to excess skew and/or kurtosis, sleep efficiency, PHQ-9 and AIS scores were transformed (see Methods). Internal reliability was acceptable to good for all psychometric questionnaires.

**Table 1.** Descriptive statistics for the main study variables. Kurtosis refers to residual kurtosis (original kurtosis-3). Note the reduction in skew and kurtosis for transformed variables which were used in subsequent analyses. AIS: Athens Insomnia Scale, TAS: Toronto Alexithymia Scale, PHQ: Patient Health Questionnaire, ZKPQ: Zuckerman-Kuhlman Personality Questionnaire, BFI: Big Five Inventory, DDF: Difficulty Describing Feelings, DIF: Difficulty Identifying Feelings, EOT: Externally Oriented Thinking.

| Variable | Mean | SD | Minimum | Maximum | Skew | Kurtosis | Cronbach's $\alpha$ | McDonald's $\Omega$ |
| --- | --- | --- | --- | --- | --- | --- | --- | --- |
| AIS | 4.59 | 3.33 | 0 | 18 | 0.98 | 0.7 | 0.779 | 0.835 |
| TAS-20 | 45.46 | 11.31 | 22 | 80 | 0.27 | -0.33 | 0.829 | 0.849 |
| DDF | 12.36 | 4.71 | 5 | 25 | 0.37 | -0.58 |  |  |
| DIF | 15.72 | 5.91 | 7 | 31 | 0.51 | -0.66 |  |  |
| EOT | 17.38 | 4.16 | 8 | 29 | 0.14 | -0.4 |  |  |
| PHQ-9 | 6.1 | 4.69 | 0 | 24 | 1.31 | 1.91 | 0.836 | 0.866 |
| Neuroticism-anxiety (ZKPQ) | 8.98 | 4.63 | 0 | 19 | 0.04 | -0.97 | 0.852 | 0.873 |
| Emotional Instability (BFI-44) | 23.84 | 6.07 | 8 | 38 | -0.32 | -0.16 | 0.836 | 0.871 |
| GSQS | 4.04 | 1.78 | 0.71 | 9.71 | 0.47 | -0.05 | 0.814 | 0.884 |
| Sleep efficiency | 90.86 | 5.73 | 37.66 | 96.95 | -4.67 | 34.79 |  |  |
| Age | 29.24 | 12.89 | 18 | 76 | 1.21 | 0.24 |  |  |
| Sleep efficiency (transformed) | 2.1 | 0.45 | 1.11 | 4.13 | 0.76 | 1.74 |  |  |
| PHQ-9 (transformed) | 1.74 | 0.7 | 0 | 3.22 | -0.44 | -0.04 |  |  |
| AIS (transformed) | 1.54 | 0.63 | 0 | 2.94 | -0.33 | -0.34 |  |  |

### Alexithymia and sleep quality

In our primary analyses, we investigated whether alexithymia (operationalized as total TAS-20 scores) is associated with subjectively assessed (GSQS, AIS) and objectively measured (EEG-based sleep efficiency) sleep quality, and if any association after controlling for negative emotionality (depressive symptoms reported on the PHQ-9, and neuroticism/emotional instability reported on the BFI-44 and ZKPQ).

After controlling for age and sex, higher TAS-20 scores were significantly associated with lower subjective sleep quality, assessed either using the GSQS (β=0.203), or the AIS (β=0.207).

This association was fully accounted for by negative emotionality. Both depressive symptoms and trait neuroticism accounted for a substantial amount of additional variance in self-reported sleep (ΔR^2^=0.067-0.296), and after the inclusion of these variables in our models, alexithymia was no longer related to subjective sleep quality.

Alexithymia was unrelated to objective sleep quality, with or without controlling for depression and personality (**Table 2**).

**Table 2.** Results from linear models regressing sleep quality on alexithymia and various controls. Separate rows show the association between alexithymia, two measures of subjective sleep quality and objective sleep quality, with or without controls for depressive affect and personality-level negative emotionality. Columns express raw (B) and standardized (β) regression coefficients for alexithymia, p-values, and variance accounted for by each model (R^2^), including incremental R^2^ (ΔR^2^) compared to the baseline model. All models are adjusted for age and sex, the effect of which are included in baseline R^2^. The sign of regression coefficients for objective sleep quality has been adjusted for the transformation of this variable, and they are shown in line with the original orientation of this variable.

| Sleep quality/model | B | $\beta$ | p | R<sup>2</sup> | $\Delta R^2$ |
| --- | --- | --- | --- | --- | --- |
| Subjective (GSQS), Raw | 0.032 | 0.203 | 0.002 | 0.049 |  |
| Subjective (GSQS), Depression-controlled | 0.007 | 0.044 | 0.486 | 0.224 | 0.175 |
| Subjective (GSQS), Personality-controlled | 0.008 | 0.048 | 0.514 | 0.13 | 0.081 |
| Subjective (AIS), Raw | 0.012 | 0.207 | 0.002 | 0.042 |  |
| Subjective (AIS), Depression-controlled | <0.001 | 0.001 | 0.982 | 0.338 | 0.296 |
| Subjective (AIS), Personality-controlled | 0.003 | 0.057 | 0.442 | 0.109 | 0.067 |
| Objective, Raw | -0.001 | -0.017 | 0.791 | 0.089 |  |
| Objective, Depression-controlled | 0.001 | 0.022 | 0.754 | 0.096 | 0.007 |
| Objective, Personality-controlled | 0.003 | 0.065 | 0.379 | 0.116 | 0.027 |

These findings are illustrated in **Figure 1**.

**Figure 1.**
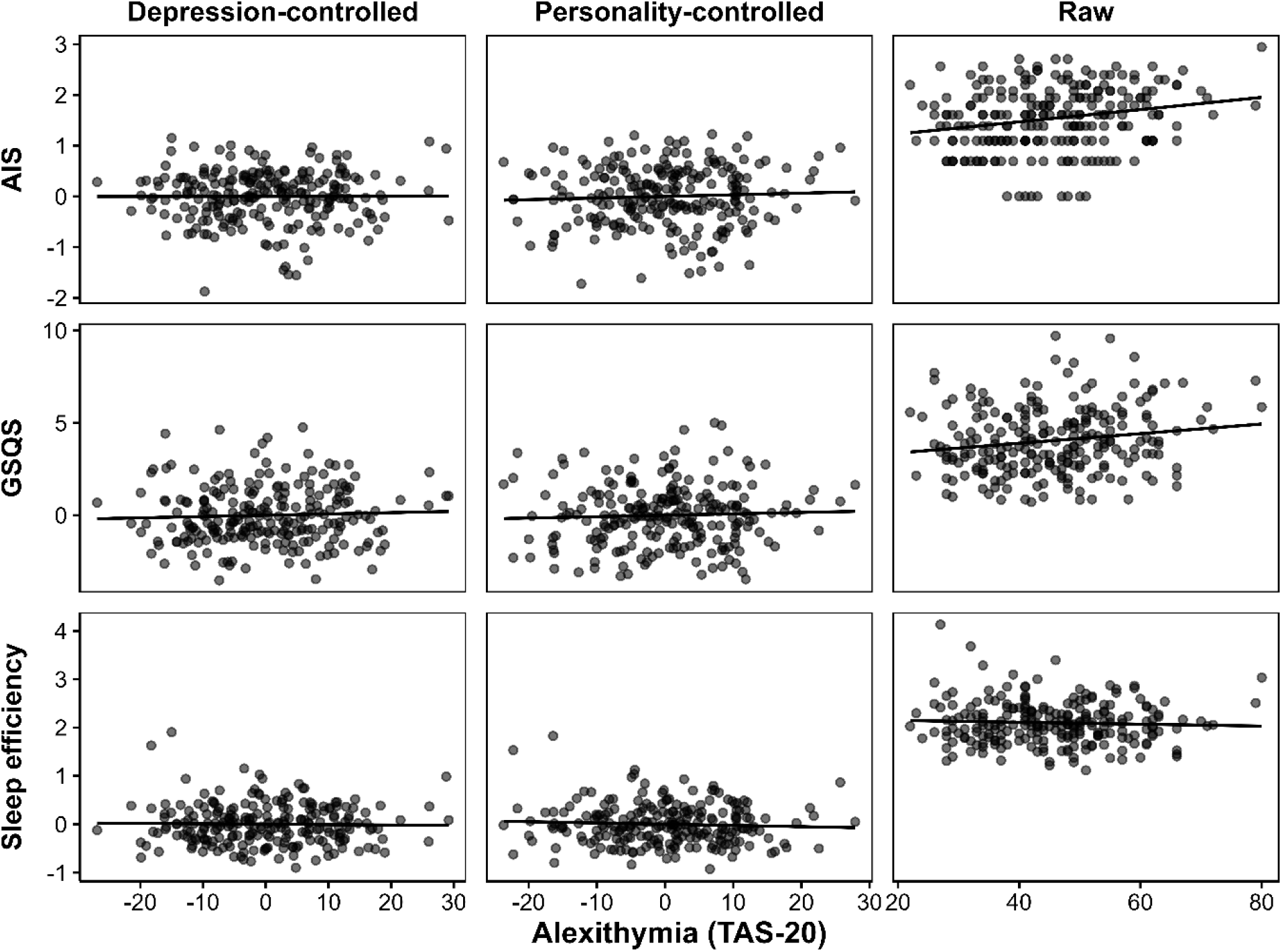
Alexithymia is only associated with subjective sleep quality, and only if psychometric confounding is not controlled. The scatterplots illustrate the raw correlations (_“_Raw_”_), and partial correlations controlling either for PHQ9 scores (_“_Depression- controlled_”_), or ZKPQ and BFI Neuroticism-anxiety/Emotional instability scores (_“_Personality-controlled_”_) between alexithymia scores on the one hand, and subjective (GSQS, AIS) and objective (Sleep efficiency) sleep quality on the other hand. Scatterplots illustrating partial correlations show zero-centered residuals. Sleep efficiency is shown after inversion and log-transformation. PHQ9 and AIS scores were also log-transformed.

### Secondary analyses

In the first set of secondary analyses, we investigated the relationship between specific alexithymia dimensions (DDF, DIF, and EOT) and both objective and subjective sleep quality. After controlling for multiple comparisons, only the association between DIF and GSQS (β=0.332, p<0.001) and AIS (β=0.353, p<0.001) was significant. After controlling for either depression or personality, the DIF-GSQS correlation lost its significance. The DIF-AIS correlation also lost its significance after controlling for depression, but it remained significant even after controlling for personality traits and multiple comparisons (β=0.233, p=0.004). These findings are shown in **Supplementary table S1**.

In the second set of secondary analyses, we investigated the relationship between objective sleep parameters other than sleep efficiency and TAS total score. No objective sleep parameter was associated with AIS scores, with or without controlling for depression and personality measures. These findings are shown in **Supplementary table S2**.

## Discussion

In this study, we investigated the link between alexithymia and the objectively and subjectively evaluated quality of sleep. We leveraged a dataset supported by objective sleep measures, multiple measures of both subjective sleep quality and possible confounders, and a sample substantially larger than any in previous investigation of objective sleep in alexithymia. We investigated three competing hypotheses about the possible mechanisms linking sleep to alexithymia: 1) overlapping physiological mechanisms, predicting changes in objective sleep quality or structure; 2) a specific alteration of subjective sleep quality, predicting no objective changes but a change in perceived sleep independent from non-specific factors; and 3) a non-specific change in subjective sleep quality only, reflecting non-specific factors such as depressive affect or neuroticism.

Our findings are at odds with the first hypothesis, as we failed to find any statistically significant link between alexithymia and either sleep efficiency (primary analysis) or any other objective sleep metric (secondary analyses). Our findings, based on a sample larger than all previous findings combined and averages of multiple nights, are therefore in line with previous negative reports (Bileviciute-Ljungar and Friberg, 2020; De Gennaro et al., 2002), and contradict an early positive finding (Bazydlo et al., 2001). Despite the appeal of this hypothesis (Godin et al., 2013), the absence of objective alterations makes it less likely that similar brain physiological mechanisms are implicated in alexithymia (a disorder of emotion processing) and sleep (a state believed to be crucial for processing emotions).

Importantly, while the findings do not support alterations in gross sleep macrostructure, it is possible that more subtle physiological changes (McCarter et al., 2022; Pierson-Bartel and Ujma, 2024) are still linked to alexithymia.

In line with a previous meta-analysis (Alimoradi et al., 2022), we found a correlation between alexithymia and poor self-reported sleep quality. In subsequent analyses, we investigated whether this is best interpreted as a specific perceived change in sleep (hypothesis 2), or the reflection of a general negative affective style, that is, a non-specific change in reported sleep quality (hypothesis 3). As the association between alexithymia and subjective sleep quality was reduced by 75-100% (**Table 2**) and no longer significant after including trait negative emotionality as a covariate, these findings contradict hypothesis 2 and support hypothesis 3 instead, suggesting that reductions of subjective sleep quality in alexithymia are non-specific. In other words, our findings suggest that people with higher alexithymia scores do not specifically perceive their sleep to be poor. Rather, their condition is associated with a negative affective style, which spills over to self-reports of sleep quality as well. Ratings of sleep are no more negative than the ratings of responders’ mood and cognitive style (depression, PHQ-9), or typical actions and characteristics (personality, ZKPQ and BFI scores).

A single exception to this trend was observed: in secondary analyses, the DIF alexithymia dimension was still significantly related to AIS scores in the personality-controlled (although not the depression-controlled) model. Additional correlations involving this variable not surviving corrections for multiple comparisons were also seen with GSQS scores, nominally significant both in the personality- (p=0.014) and depression-controlled (p=0.031) models.

These findings suggest that interoceptive symptoms of alexithymia – the ability to assess which emotions physiological states correspond to – may interfere with assessing subjective sleep quality specifically. Thus, in the context of the Difficulty Identifying Symptoms dimension, hypothesis 2 receives partial support: sleep quality may be systematically perceived as worse in those who struggle with labeling internal physiological states. While this finding is not unambiguously supported by both subjective sleep metrics and both measures of subjective sleep quality, it is theoretically feasible and warrants further investigation.

Our work has limitations. First, we assessed alexithymia symptoms in a healthy sample. Our findings may not generalize to sleep alterations in individuals with clinically significant levels of alexithymia. We note, however, that almost all research in this field has been similarly conducted in similar samples because, as alexithymia is not currently recognized as a separate psychiatric condition, the collection of a true patient sample is challenging.

Second, while our EEG system has been validated against the polysomnography gold standard, it is not PSG itself. Third, our findings about the absence of objective sleep EEG alterations in alexithymia may not preclude the absence of any such alterations at all. While our investigation explored the most frequently used gross macrostructural variables and found no alterations, it is possible that alexithymia does affect more fine-grained aspects of objectively measured sleep.

In conclusion, our work found that alexithymia is associated only with subjective alterations of sleep, and these mostly reflect a non-specific effect of elevated negative emotionality rather than a specific change in perceived sleep quality. Our findings make it less likely that similar physiological mechanisms are implicated in sleep-related emotional processing and alexithymia and suggest that the sleep of individuals with clinically significant levels of alexithymia may benefit most from interventions improving non-specific affective complaints rather than the core symptomatology of this condition related to processing emotions. A possible exception, however, is the difficulty of labeling interoceptive experiences (DIF), which appears specifically related to poor sleep quality.

## Author’s contribution

Pardis Adibi: Conceptualization, Formal analysis, Investigation, Data Curation, Visualization, Writing – original draft Péter Przemyslaw Ujma: Conceptualization, Methodology, Resources, Supervision, Writing – review and editing

## Funder involvement

The funders had no role in the design of the study, data collection, analysis, interpretation, or in the writing of the manuscript.

## References

Alimoradi, Z., Majd, N.R., Broström, A., Tsang, H.W.H., Singh, P., Ohayon, M.M., Lin, C.-Y., Pakpour, A.H., 2022. Is alexithymia associated with sleep problems? A systematic review and meta-analysis. Neurosci. Biobehav. Rev. 133, 104513. 10.1016/j.neubiorev.2021.12.036.

Arnal, P.J., Thorey, V., Debellemaniere, E., Ballard, M.E., Bou Hernandez, A., Guillot, A., Jourde, H., Harris, M., Guillard, M., Van Beers, P., Chennaoui, M., Sauvet, F., 2020. The Dreem Headband compared to polysomnography for electroencephalographic signal acquisition and sleep staging. Sleep 43. 10.1093/sleep/zsaa097.

Bagby, R.M., Parker, J.D., Taylor, G.J., 1994. The twenty-item Toronto Alexithymia Scale--I. Item selection and cross-validation of the factor structure. J. Psychosom. Res. 38, 23–32. 10.1016/0022-3999(94)90005-1.

Bazydlo, R., Lumley, M.A., Roehrs, T., 2001. Alexithymia and polysomnographic measures of sleep in healthy adults. Psychosom. Med. 63, 56–61. 10.1097/00006842-200101000-00007.

Bileviciute-Ljungar, I., Friberg, D., 2020. Emotional Awareness Correlated With Number of Awakenings From Polysomnography in Patients With Myalgic Encephalomyelitis/Chronic Fatigue Syndrome-A Pilot Study. Front. Psychiatry 11, 222. 10.3389/fpsyt.2020.00222.

Catherman, C., Cassidy, S., Benca-Bachman, C.E., Barber, J.M., Palmer, R.H.C., 2023. Associations between neuroticism, subjective sleep quality, and depressive symptoms across the first year of college. J. Am. Coll. Health 71, 381–388. 10.1080/07448481.2021.1891917.

Cserjési, R., Luminet, O., Lénárd, L., 2007. A Torontói Alexitímia Skála (TAS-20) magyar változata: megbízhatósága és faktorvaliditása egyetemista mintán. Magyar Pszichológiai Szemle 62, 355–368. 10.1556/mpszle.62.2007.3.4.

De Gennaro, L., Ferrara, M., Curcio, G., Cristiani, R., Lombardo, C., Bertini, M., 2002. Are polysomnographic measures of sleep correlated to alexithymia? A study on laboratory-adapted sleepers. J. Psychosom. Res. 53, 1091–1095. 10.1016/s0022-3999(02)00342-2.

De Gennaro, L., Martina, M., Curcio, G., Ferrara, M., 2004. The relationship between alexithymia, depression, and sleep complaints. Psychiatry Res. 128, 253–258. 10.1016/j.psychres.2004.05.023.

Dreem Inc, 2017. Dreem Whitepaper. Dreem Inc.

Godin, I., Montplaisir, J., Gagnon, J.-F., Nielsen, T., 2013. Alexithymia associated with nightmare distress in idiopathic REM sleep behavior disorder. Sleep 36, 1957–1962. 10.5665/sleep.3238.

Han, S.-H., Choo, Y.-S., Koo, G.E., Kang, Y.-J., 2023. Is alexithymia associated with sleep disturbances, independent of depression and anxiety? J. Sleep Med. 20, 41–46. 10.13078/jsm.230006.

Hendryx, M.S., Haviland, M.G., Shaw, D.G., 1991. Dimensions of alexithymia and their relationships to anxiety and depression. J. Pers. Assess. 56, 227–237. 10.1207/s15327752jpa5602_4.

Hogeveen, J., Grafman, J., 2021. Alexithymia. Handb. Clin. Neurol. 183, 47–62. 10.1016/B978-0-12-822290-4.00004-9.

Kövi, Z., Rózsa, S., Takács, M., Takács, S., Hevesi, K., Vargha, A., 2018. ZUCKERMAN-KUHLMAN-ALUJA SZEMÉLYISÉGKÉRDŐíV (ZKA-PQ) MAGYAR VERZIÓJA. PSYHUNG.

Kroenke, K., Spitzer, R.L., Williams, J.B., 2001. The PHQ-9: validity of a brief depression severity measure. J. Gen. Intern. Med. 16, 606–613. 10.1046/j.1525-1497.2001.016009606.x.

Marchesi, C., Brusamonti, E., Maggini, C., 2000. Are alexithymia, depression, and anxiety distinct constructs in affective disorders? J. Psychosom. Res. 49, 43–49. 10.1016/s0022-3999(00)00084-2.

Ma, Q., Zhang, X., Zou, L., 2020. The mediating effect of alexithymia on the relationship between schizotypal traits and sleep problems among college students. Front. Psychiatry 11, 153. 10.3389/fpsyt.2020.00153.

McCarter, S.J., Hagen, P.T., St Louis, E.K., Rieck, T.M., Haider, C.R., Holmes, D.R., Morgenthaler, T.I., 2022. Physiological markers of sleep quality: A scoping review. Sleep Med. Rev. 64, 101657. 10.1016/j.smrv.2022.101657.

Pierson-Bartel, R., Ujma, P.P., 2024. Objective sleep quality predicts subjective sleep ratings. Sci. Rep. 14, 5943. 10.1038/s41598-024-56668-0.

Ricciardi, L., Demartini, B., Fotopoulou, A., Edwards, M.J., 2015. Alexithymia in neurological disease: A review. J. Neuropsychiatry Clin. Neurosci. 27, 179–187. 10.1176/appi.neuropsych.14070169.

Ronai, K.Z., Szentkiralyi, A., Lazar, A.S., Lazar, Z.I., Papp, I., Gombos, F., Zoller, R., Czira, M.E., Lindner, A.V., Mucsi, I., Bodizs, R., Molnar, M.Z., Novak, M., 2017. Association of symptoms of insomnia and sleep parameters among kidney transplant recipients. J. Psychosom. Res. 99, 95–104. 10.1016/j.jpsychores.2017.05.019.

Rózsa, S., 2010. A Big Five Inventory magyar adaptációja. Eötvös Loránd Tudományegyetem, Budapest.

Sifneos, P.E., 1973. The prevalence of “alexithymic” characteristics in psychosomatic patients. Psychother. Psychosom. 22, 255–262. 10.1159/000286529.

Simor, P., Köteles, F., Bódizs, R., Bárdos, G., 2009. A questionnaire based study of subjective sleep quality: The psychometric evaluation of the Hungarian version of the Groningen Sleep Quality Scale. Mentálhigiéné és Pszichoszomatika 10, 249–261. 10.1556/Mental.10.2009.3.5.

Svetnik, V., Snyder, E.S., Tao, P., Roth, T., Lines, C., Herring, W.J., 2020. How well can a large number of polysomnography sleep measures predict subjective sleep quality in insomnia patients? Sleep Med. 67, 137–146. 10.1016/j.sleep.2019.08.020.

Taji, W., Pierson, R., Ujma, P.P., 2023. Protocol of the Budapest sleep, experiences, and traits study: An accessible resource for understanding associations between daily experiences, individual differences, and objectively measured sleep. PLoS ONE 18, e0288909. 10.1371/journal.pone.0288909.

Taylor, G.J., Bagby, R.Michael., Parker, J.D.A. (James D.A., 1999. Disorders of affect regulation: Alexithymia in medical and psychiatric illness. Cambridge.

Utsumi, T., Yoshiike, T., Kaneita, Y., Aritake-Okada, S., Matsui, K., Nagao, K., Saitoh, K., Otsuki, R., Shigeta, M., Suzuki, M., Kuriyama, K., 2022. The association between subjective-objective discrepancies in sleep duration and mortality in older men. Sci. Rep. 12, 18650. 10.1038/s41598-022-22065-8.

