## Supplemental Tables and Figures for "Sleep complaints in alexithymia reflect non-specific negative affectivity"

| Sleep Quality/Model | Predictor | B | Beta | p | $R^2$ | $\Delta R^2$ |
|---|---|---|---|---|---|---|
| GSQS, Raw | DDF | 0.047 | 0.125 | 0.066 | 0.024 | |
| GSQS, Depression-controlled | DDF | -0.009 | -0.024 | 0.707 | 0.222 | 0.198 |
| GSQS, Personality-controlled | DDF | -0.001 | -0.003 | 0.964 | 0.128 | 0.104 |
| GSQS, Raw | DIF | 0.1 | 0.332 | <0.001 | 0.11 | |
| GSQS, Depression-controlled | DIF | 0.044 | 0.147 | 0.031 | 0.238 | 0.127 |
| GSQS, Personality-controlled | DIF | 0.059 | 0.196 | 0.014 | 0.152 | 0.047 |
| GSQS, Raw | EOT | -0.014 | -0.033 | 0.624 | 0.01 | |
| GSQS, Depression-controlled | EOT | -0.013 | -0.029 | 0.625 | 0.223 | 0.213 |
| GSQS, Personality-controlled | EOT | -0.031 | -0.073 | 0.25 | 0.134 | 0.124 |
| AIS, Raw | DDF | 0.023 | 0.168 | 0.013 | 0.027 | |
| AIS, Depression-controlled | DDF | -0.002 | -0.017 | 0.767 | 0.338 | 0.311 |
| AIS, Personality-controlled | DDF | 0.007 | 0.049 | 0.488 | 0.108 | 0.081 |
| AIS, Raw | DIF | 0.038 | 0.353 | <0.001 | 0.114 | |
| AIS, Depression-controlled | DIF | 0.011 | 0.105 | 0.095 | 0.346 | 0.232 |
| AIS, Personality-controlled | DIF | 0.025 | 0.233 | 0.004 | 0.139 | 0.025 |
| AIS, Raw | EOT | -0.015 | -0.098 | 0.147 | 0.009 | |
| AIS, Depression-controlled | EOT | -0.014 | -0.093 | 0.09 | 0.346 | 0.337 |
| AIS, Personality-controlled | EOT | -0.021 | -0.136 | 0.034 | 0.124 | 0.113 |
| Objective, Raw | DDF | -0.003 | -0.033 | 0.614 | 0.09 | |
| Objective, Depression-controlled | DDF | -0.007 | -0.072 | 0.292 | 0.1 | 0.01 |
| Objective, Personality-controlled | DDF | -0.009 | -0.095 | 0.176 | 0.121 | 0.032 |
| Objective, Raw | DIF | 0.007 | 0.095 | 0.154 | 0.097 | |
| Objective, Depression-controlled | DIF | 0.004 | 0.059 | 0.424 | 0.098 | 0.01 |
| Objective, Personality-controlled | DIF | 0.002 | 0.022 | 0.787 | 0.114 | 0.015 |
| Objective, Raw | EOT | -0.005 | -0.044 | 0.499 | 0.091 | |
| Objective, Depression-controlled | EOT | -0.005 | -0.043 | 0.506 | 0.097 | 0.006 |
| Objective, Personality-controlled | EOT | -0.007 | -0.063 | 0.323 | 0.117 | 0.026 |

**Supplementary table S1.** Results from linear models regressing sleep quality on alexithymia dimensions. Separate rows show the association between alexithymia dimensions DDF, DIF and EOT, two measures of subjective sleep quality and objective sleep quality, with our without controls for depressive affect and personality-level negative emotionality. Columns express raw (B) and standardized (Beta) regression coefficients for alexithymia, p-values, and variance accounted for by each model ($R^2$), including incremental $R^2$ ($\Delta R^2$) compared to the baseline model. All models are adjusted for age and sex.

| Sleep parameter/model | B | Beta | p | $R^2$ | $\Delta R^2$ |
|---|---|---|---|---|---|
| SOL, Raw | 0 | -0.01 | 0.875 | -0.007 | |
| SOL, Depression-controlled | -0.001 | -0.027 | 0.706 | -0.008 | -0.001 |
| SOL, Personality-controlled | -0.001 | -0.02 | 0.798 | 0.004 | 0.011 |
| WASO, Raw | 0.002 | 0.037 | 0.536 | 0.158 | |
| WASO, Depression-controlled | 0 | 0.004 | 0.95 | 0.161 | 0.003 |
| WASO, Personality-controlled | 0 | -0.008 | 0.912 | 0.151 | -0.007 |
| TST, Raw | -0.152 | -0.029 | 0.653 | -0.003 | |
| TST, Depression-controlled | 0.145 | 0.028 | 0.688 | 0.012 | 0.015 |
| TST, Personality-controlled | 0.335 | 0.065 | 0.393 | 0.012 | 0.015 |
| N1 percentage, Raw | -0.01 | -0.055 | 0.37 | 0.117 | |
| N1 percentage, Depression-controlled | -0.023 | -0.129 | 0.047 | 0.147 | 0.03 |
| N1 percentage, Personality-controlled | -0.018 | -0.1 | 0.164 | 0.131 | 0.014 |
| N2 percentage, Raw | -0.028 | -0.045 | 0.436 | 0.2 | |
| N2 percentage, Depression-controlled | -0.011 | -0.017 | 0.781 | 0.206 | 0.006 |
| N2 percentage, Personality-controlled | 0.019 | 0.031 | 0.655 | 0.21 | 0.001 |
| N3 percentage, Raw | 0.032 | 0.049 | 0.388 | 0.245 | |
| N3 percentage, Depression-controlled | 0.029 | 0.044 | 0.47 | 0.243 | -0.002 |
| N3 percentage, Personality-controlled | 0.014 | 0.022 | 0.746 | 0.245 | 0 |
| REM percentage, Raw | 0.006 | 0.014 | 0.829 | -0.004 | |
| REM percentage, Depression-controlled | 0.005 | 0.012 | 0.865 | -0.009 | -0.005 |
| REM percentage, Personality-controlled | -0.015 | -0.035 | 0.651 | -0.004 | 0 |
| REM latency, Raw | 0.001 | 0.023 | 0.726 | 0.023 | |
| REM latency, Depression-controlled | 0 | 0.017 | 0.809 | 0.02 | -0.003 |
| REM latency, Personality-controlled | 0.001 | 0.051 | 0.504 | 0.018 | -0.005 |
| Awakenings, Raw | -0.006 | -0.009 | 0.893 | 0.022 | |
| Awakenings, Depression-controlled | -0.04 | -0.062 | 0.371 | 0.033 | 0.011 |
| Awakenings, Personality-controlled | -0.017 | -0.025 | 0.742 | 0.015 | -0.007 |

**Supplementary table S2.** Results from linear models regressing additional objective sleep parameters on alexithymia and various controls. Separate rows show the association between alexithymia and sleep parameters with our without controls for depressive affect and personality-level negative emotionality. Columns express raw (B) and standardized (Beta) regression coefficients for alexithymia, p-values, and variance accounted for by each model ($R^2$), including incremental $R^2$ ($\Delta R^2$) compared to the baseline model. All models are adjusted for age and sex.

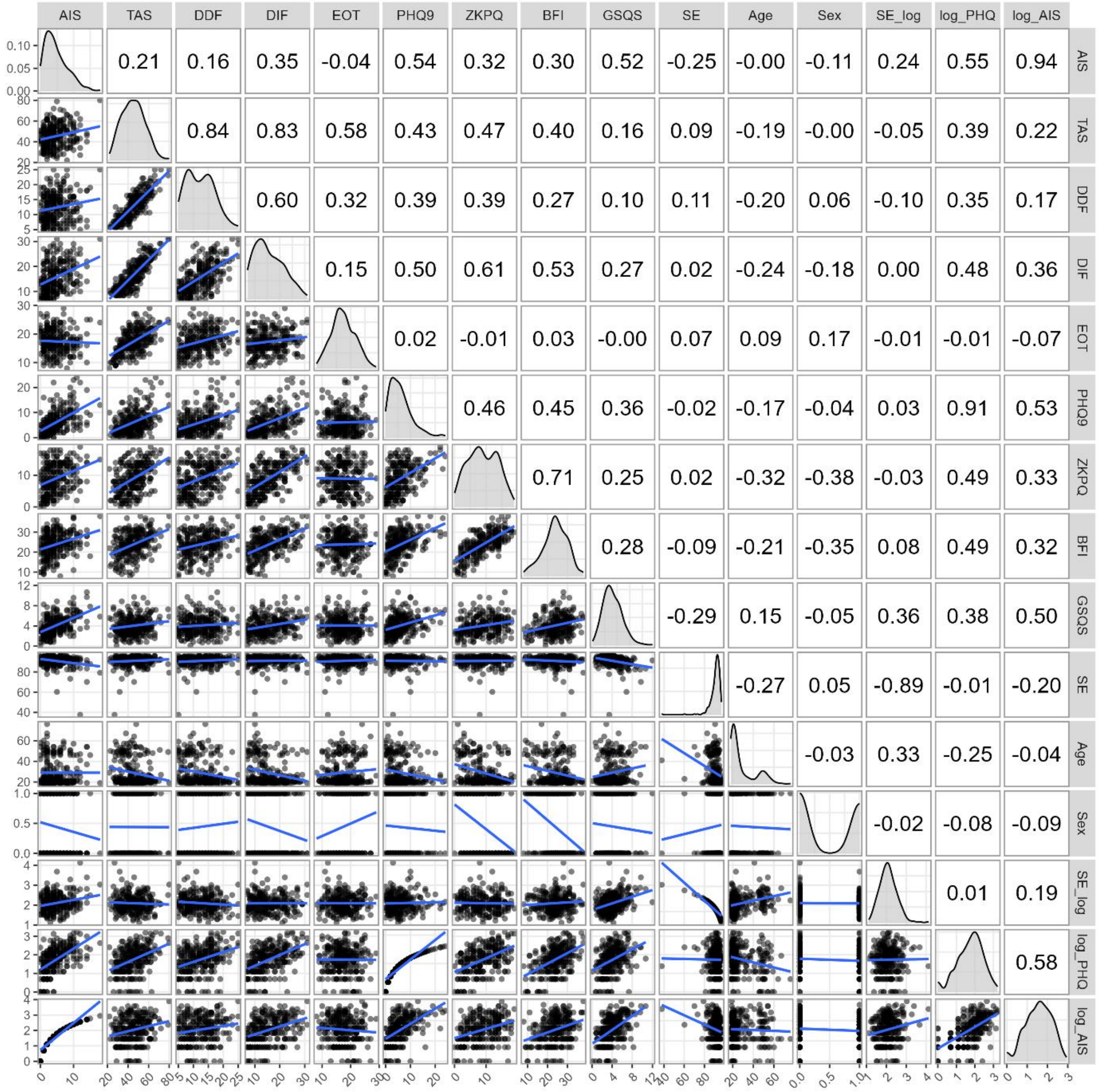


**Supplementary Figure S1**. Kernel density histograms (diagonals), scatterplots (lower triangle) and correlation coefficients (upper triangle) of key variables. For variables which required log-transforming, both the original and the transformed version is displayed. AIS: Athens Insomnia Scale, TAS: Toronto Alexithymia Scale total score, DDF: Difficulty Describing Feelings, DIF: Difficulty Identifying Feelings, EOT: Externally Oriented Thinking, PHQ9: PHQ-9 total score, ZKPQ: ZKPQ Neuroticism-Anxiety score, BFI: BFI Emotional Instability score, GSQS: Groningen Sleep Quality Scale mean score, SE: Sleep Efficiency. For the binary variable Sex, 0 refers to females and 1 to males.
